# SeqPIP-2020: Sequence based Protein Interaction Prediction Contest

**DOI:** 10.1101/2020.11.12.380774

**Authors:** Anup Kumar Halder, Ayatullah Faruk Mollah, Piyali Chatterjee, Dipak Kumar Kole, Subhadip Basu, Dariusz Plewczyński

## Abstract

Computational protein-protein interaction (PPI) prediction techniques can contribute greatly in reducing time, cost and false-positive interactions compared to experimental approaches. Sequence is one of the key and primary information of proteins that plays a crucial role in PPI prediction. Several machine learning approaches have been applied to exploit the characteristics of PPI datasets. However, these datasets greatly influence the performance of predicting models. So, care should be taken on both dataset curation as well as design of predictive models. Here, we summarize the results of the SeqPIP competition whose objective was to develop comprehensive PPI predictive models from sequence information with high-quality bias-free interaction datasets. A training set of 2000 positive and 2000 negative interactions with sequences was given to each contestant. The methods were evaluated with three independent high-quality interaction test datasets.

## 1 Introduction

Protein works by interacting with other proteins to accomplish various biological processes essential to a living organism. Identification of such interactions facilitates different biological problems such as therapeutic target identification [10], characterization of metabolic and cellular activity, hormone regulation, signal transduction, DNA transcription and replication, exploring the pathogenesis of disease [4], drug target identification [7] etc. Protein-protein interactions (PPIs) detection has been improved with high throughput experimental technologies but on a small portion. These experimental approaches are expensive and time-consuming to apply over large-scale PPIs. Thus, there is immense need for reliable computational approaches to identify and characterize PPIs. The primary structure of a protein, represented by amino acid sequences is the simplest type of information that can be applied in PPI prediction through intuitive numerical feature representation. There are plenty of information in primary sequence of a protein and experimental studies reveal that the sequence is one of the key information for PPI prediction [14,11,13,6, 12]. The experimental studies become the evidence of applicability and efficacy of primary structure data in resolving different complex proteomic properties.

In recent studies [8,5,9], it has been observed that in most of the cases the PPI datasets used for evaluation were biased due to *component level overlapping.* For any PPI pair, sharing of individual component (sequence information) between training and test sets is referred to as *component-level overlapping*. For an unbiased and fair evaluation of the PPI predictive model, it is necessary to distinguish the test pairs on the basis of whether they share individual sequence with any pair of the training set [8]. The frequent occurrences of individual protein and/or pair in both training and test data raises the component level overlapping issue and bias the model greatly. To avoid this bias in PPI prediction, a dataset curation scheme is proposed by [8, 5] where three levels of complex test classes are introduced for the predictive models. Park *et al.* [8] experimentally established that the performance of the top methods on unbiased datasets is significantly lower than their previously published results which raises the bar higher for PPI prediction methods. In this competition, we have designed and curated the dataset by removing the component-level overlapping issue to make the PPI prediction unbiased and comprehensive.

We organized the *Seq*uence based *P*rotein *I*nteraction *P*rediction competition *(SeqPIP*) at International Conference on Frontiers in Computing and Systems (COMSYS-2020) to build upon the high quality PPI dataset and encourage researchers across the world to introduce new techniques for sequence based generalized and unbiased PPI prediction. In sequence based PPI prediction, the main computational challenge is to find suitable way to describe important feature information. Moreover, dataset preparation is very crucial for machine learning methods. Here, unbiased training and test datasets are carefully curated and the competition rules are concocted in accordance with these objectives.

Through this competition, several sequence based approaches have been introduced to improve the performances on the provided benchmark dataset. In this paper, we describe in detail the dataset and the emerging trends of techniques that performed well on the aforesaid challenging task. We hope that the algorithms described in this competition will be of use to the broader community of bioinformatics research. The paper is organized as follows. We describe the unbiased dataset creation and its properties along with the competition rules in Section 2. Then, a summary of methods developed by the participants are presented in Section 3. In Section 4, comparative assessment with standard metrics and related discussion are presented. Finally, conclusion is made in Section 5.

## 2 Dataset Preperation

We have retrieved the positive interaction data from HIPPIE (v2: 1) [2] database and high quality subset of these datasets is selected for the competition. The high quality PPI data is selected based on confidence scoring (larger than 0.8) of HIPPIE. The negative protein-protein interaction (NPPI) data is generated by random sampling of protein pairs that are not known to interact in any benchmark PPI dataset. The sequence information is retrieved from UniprotKB/SwissProt database [3]. Protein sequences which are too similar should not occur simultaneously in training and test sets. Presence of redundant homologous sequences in PPI dataset greatly influences the performance of predictive models. To remove these homologous sequences, CD-Hit [1] is applied over the unique sequences that are present in the dataset with a threshold of 0.4 (40% identity). This ensures that, there is no issue of occurrences for identical sequences (≤ 40%) that reside either between train/test or within train/test as pairs. Total 5435 unique non-redundant protein sequences are retrieved for final PPI dataset selection.

Finally, 4500 positive and 4500 negative interaction pairs are selected from previously extracted high confidence PPIs and randomly generated NPPIs respectively. All 9000 interactions are comprised by 5435 non-redundant unique sequences. To make the competition comprehensive and unbiased, the dataset is curated carefully and partitioned into training and test sets by removing component level overlapping as proposed by [8]. To implement the idea of removing *component-level overlapping* issue, test data is designed into three difficulty classes viz. C1, C2 and C3. The unbiased data preparation and detailed description of all three test classes (C1, C2 and C3) are discussed below.

We have implemented a scheme similar to what was proposed by Park and Marcotte [8] to design the unbiased dataset for sequence based PPI prediction. For an unbiased and fair evaluation of the PPI predictive model, it is necessary to distinguish the test pairs on the basis of whether they share similar sequence information with the pairs of the train set. Here, the similarity is considered as over 40% identity between any two sequences. To overcome these issues, Park and Marcotte [8] proposed a scheme that, for any trainset the test cases will be partitioned into three distinct predictive test classes (C1, C2 and C3). In C1, both sequences of any test pair may be present in the trainset but not as a pair. In C2, only one component (sequence) can be present in the trainset and in C3, no components in the test pair could be present in the trainset. Detailed dataset curation and train/test partitioning is demonstrated with a toy example (with Graph based representation) in Fig. 1.

**Fig. 1.**
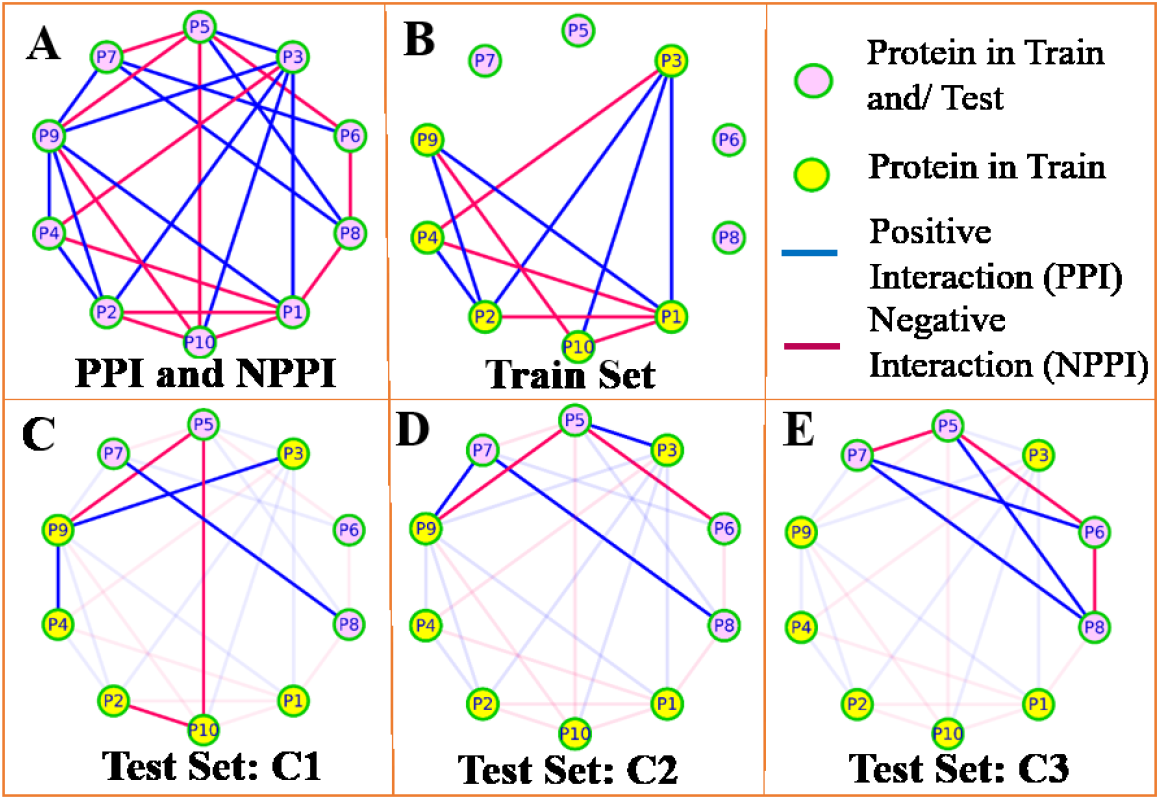
Schematic diagram of data preparation. A) protein interaction network with PPI (red edges) and NPPI (blue edges). B) Selected Train set with the set of nodes (P1, P2, P3, P4, P9, P10), All the nodes in trainset are marked with yellow color (from B to E). C) Test set C1, with the set of nodes (P2, P3, P4, P9, P10), D) Test set C2, with the set of nodes (P3, P5, P6, P7, P8, P9). E) Test set C3, with the set of nodes (P5, P6, P7, P8). All three test classes ensure that they have not shared any exact pair (same edge) with the trainset. C1 has component level overlapping as well as pair (both components) sharing as both train and C1 shares P3, P2, P4, P9 and P10. For example, both proteins P3 and P9 from pair P3-P9 in C1 is also present in the trainset, and similarly for pair P2-P10, P4-P9. In C2, only one protein node is shared between train and test set. For example, in pair, P9-P5, P3-P5 only one node from each pair is shared such as P9, P3 respectively. In C3, no edges and nodes are shared between train and test set.

Total 4000 (2000 positive and 2000 negative) interactions were provided to the competitors as trainset. However, the test datasets were not shared to the competitors before announcement of results and the competitors’ models were evaluated only once on the test datasets after the final submission of their models. Details of the competition datasets are shown in Table 1.

**Table 1:** SeqPIP-2020 competition dataset details (C1, C2 and C3 are the three subsets of testset).

| Dataset Type | Total Interactions | Positive (PPI) | Negative (NPPI) | # of Unique Sequences |
| --- | --- | --- | --- | --- |
| *Train* | 4000 | 2000 | 2000 | 3374 |
| *C1* | 2000 | 1000 | 1000 | 1672 |
| *C2* | 1500 | 750 | 750 | 978 |
| *C3* | 1500 | 750 | 750 | 984 |
| *Total* | 9000 | 4500 | 4500 | 5435 |

## 3 Participating Methods

Here, we present the best four methods submitted to SeqPIP. Among these methods, two are based on machine learning and the other two are based on sequence similarity. In the following sections, we will denote the machine learning based methods as ML1 and ML2, and sequence similarity based approaches as SqS1 and SqS2 while the ordering of methods is determined by evaluation performance.

### 3.1 ML1: *Soumyadeep Debnath, Tata Consultancy Services and Ayatullah Faruk Mollah, Aliah University, India*

The feature vector was extracted for every protein pair from the dataset. The sequence information is represented by 12 physicochemical properties of amino acids, namely, hydrophilicity (H2), flexibility (F), accessibility (A1), turns scale (T), exposed surface (E), polarity (Pa), antigenic propensity (A2), hydrophobicity (Ha), net charge index of the side chains (NCI), polarizability (P2), solvent accessible surface area (SASA), and side-chain volume (V). Among these 12 properties, hydrophobicity and polarity are calculated according to two different scales or methods like H11a, H12a and P11a, P12a respectively. Based on the values of these 14 physicochemical property scales of 20 essential amino acids, 14-length vectors are extracted from all amino acids for every protein sequence. For any pair of protein sequences, the final data is generated by computing the mean value of the two 14-length sequence feature vectors. With this feature representation of the train dataset, the model is trained by linear kernel driven support vector machine (SVM).

### 3.2 ML2: *Kaustav Sengupta, University of Warsaw, Poland*

In this method, two level approach has been used for PPI prediction, i) SVM based feature selection and ii) random forest based PPI prediction on the selected features. Feature selection method is carried out over 554 amino acid indices (AAI) and resulted with the best 22 AAI features. Then, the method has extracted the feature values for both the protein sequences using selected 22 AAI indices and concatenated for final data representation. This sequence based information is applied to random forest (number of estimator = 150, max depth = 8, min sample leaf = 200) classifier for final model generation.

### 3.3 SqS1: *Dipayan Ray, Dr. Sudhir Chandra Sur Degree Engineering College, India*

In SqS1, sequence similarity based approach is proposed where it intuitively explores the sequence information in PPI detection. For each sequence, n-gram token set is generated where exploring the n values up to 5. Then the similarity between the sequence pair is defined as the Jaccard similarity of the generated tokens. A predefined threshold is identified from the train dataset at which the accuracy become high and based on this cutoff score the interactions are classified as positive and negative in the test datasets of the competitions.

### 3.4 SqS2: *Rohan Sarkar, Dr. Sudhir Chandra Sur Degree Engineering College, India*

SqS2 is one of the non-machine learning (ML) based approach on SeqPIP. Here, Jaro–Winkler distance (JW) is introduced to define the similarity between the sequence pairs. For any pair of sequences, the higher JW distance indicates more sequence similarity and ranges from 0 (no similarity) to 1 (exact matching). In this approach, higher sequence similarity between the sequences are considered as better candidate for the positive interactions. A cutoff score is detected by the method at which it maximise the accuracy on the train dataset. Finally, with this similarity metric and threshold value, all three test datasets are evaluated and reported.

## 4 Results and Discussion

We have compared performance of the best methods of SeqPIP. All the models are tested on the test datasets and the results are reported in Table 2. Different statistical metrics such as precision, recall, accuracy and F1-score are computed to evaluate the models. The results in Table 2 suggest that *ML1* has achieved better performance in test classes C1 and C3 where *ML2* has highest recall (0.631) and accuracy (0.623) in C2 test class. Among the non-ML based approaches, *SqS1* has performed better than *SqS2.* Although, the non-ML based approaches are not comprehensive in the performance evaluation metric compared to ML based methods.

**Table 2.**
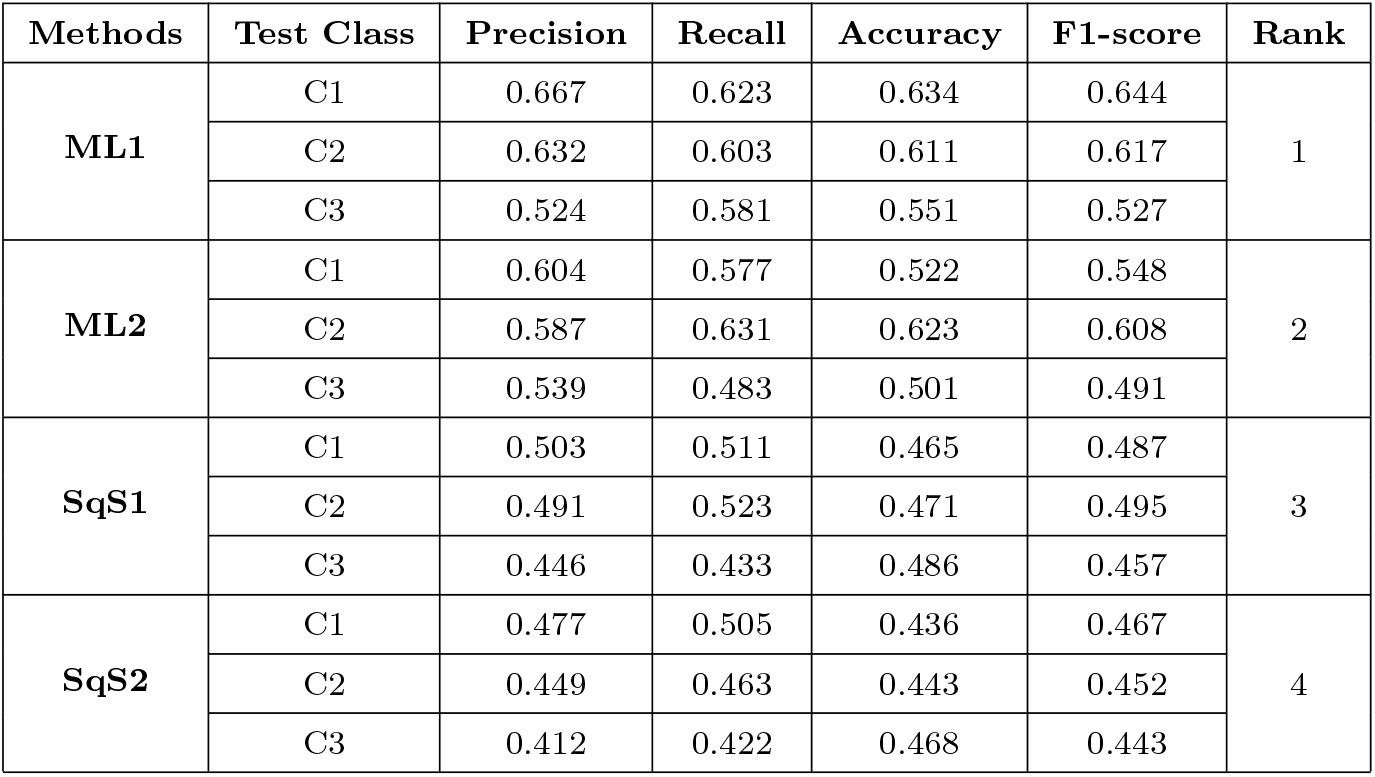
Performance evaluation on best 4 methods of SeqPIP.

| Methods | Test Class | Precision | Recall | Accuracy | F1-score | Rank |
| --- | --- | --- | --- | --- | --- | --- |
| **ML1** | C1 | 0.667 | 0.623 | 0.634 | 0.644 | 1 |
|  | C2 | 0.632 | 0.603 | 0.611 | 0.617 |  |
|  | C3 | 0.524 | 0.581 | 0.551 | 0.527 |  |
| **ML2** | C1 | 0.604 | 0.577 | 0.522 | 0.548 | 2 |
|  | C2 | 0.587 | 0.631 | 0.623 | 0.608 |  |
|  | C3 | 0.539 | 0.483 | 0.501 | 0.491 |  |
| **SqS1** | C1 | 0.503 | 0.511 | 0.465 | 0.487 | 3 |
|  | C2 | 0.491 | 0.523 | 0.471 | 0.495 |  |
|  | C3 | 0.446 | 0.433 | 0.486 | 0.457 |  |
| **SqS2** | C1 | 0.477 | 0.505 | 0.436 | 0.467 | 4 |
|  | C2 | 0.449 | 0.463 | 0.443 | 0.452 |  |
|  | C3 | 0.412 | 0.422 | 0.468 | 0.443 |  |

The training set and the test sets at three component levels viz. C1, C2 and C3 have been made publicly available for academic, research and noncommercial purposes at https://sites.google.com/site/bioinfoju/home/seqpip

## 5 Conclusion

Here, we have presented the sequence based PPI prediction approaches and their performances in connection with the SeqPIP-2020 competition. Different machine learning based and non-machine learning based approaches have been employed to predict the PPIs by exploring the primary sequence information of proteins. All the methods are trained and evaluated on unbiased train and test data sets by removing *component level overlapping* biases. Among the best four approaches, the overall performance of machine learning based approaches are found superior than other approaches. It may be stated that the datasets and the methods of SeqPIP enable a platform for many bioinformatics applications such as complex detection, characterisation of functional relationship, PPI network analysis, etc. The datasets are also made available as a reference for research purposes.

